# The Clinical and Molecular Epidemiology of CTX-M-9 Group Producing Enterobacteriaceae infections in children

**DOI:** 10.1101/416016

**Authors:** Latania K. Logan, Rachel L. Medernach, T. Nicholas Domitrovic, Jared R. Rispens, Andrea M. Hujer, Nadia K. Qureshi, Steven H. Marshall, David C. Nguyen, Susan D. Rudin, Xiaotian Zheng, Robert A. Weinstein, Robert A. Bonomo

## Abstract

**Background:** The pandemic of extended-spectrum-beta-lactamase (ESBL)-producing-Enterobacteriaceae (Ent) is strongly linked to the dissemination of CTX-M-type-ESBL-Ent. We sought to define the epidemiology of infections in children due to an emerging resistance type, CTX-M-9-group-producing-Ent (CTX-M-9-grp-Ent).

**Methods:** A multi-centered case-control analysis of Chicago children with CTX-M-9-grp-Ent infections was performed. Cases were defined as children possessing extended-spectrum-cephalosporin-resistant (ESC-R) infections. PCR and DNA analysis assessed beta-lactamase (*bla*) genes, multi-locus sequence types (MLST) and phylogenetic grouping of *E. coli*. Controls were children with ESC-susceptible (ESC-S)-Ent infections matched 3:1 by age, source, and hospital. The clinical-epidemiologic predictors of CTX-M-9-grp-Ent infection were assessed.

**Results:** Of 356 ESC-R-Ent isolates from children (median age 4.1 years), CTX-M-9-group was the solely detected *bla* gene in 44(12.4%). The predominant species was *E. coli* (91%) of virulent phylogroups D(60%) and B2(40%). MLST revealed multiple strain types. On multivariable analysis, CTX-M-9-grp-Ent occurred more often in *E. coli* (OR 7.0), children of non-black-white-Hispanic race (OR 6.5), and outpatients (OR 4.5) which was a very unexpected finding for infections due to antibiotic-resistant bacteria. Residents of South Chicago were 6.7 times more likely to have CTX-M-9-grp-Ent infections than those in the reference region (West), while residence in Northwestern Chicago was associated with an 81% decreased risk. Other demographic, comorbidity, invasive-device, and antibiotic use differences were not found.

**Conclusions:** CTX-M-9-grp-Ent infection is strikingly associated with patient residence and is occurring in children without traditional in-patient exposure risk factors. This suggests that among children, the community environment may be a key contributor in the spread of these resistant pathogens.

## Introduction

The pandemic of multi-drug resistant (MDR) Enterobacteriaceae remains one of the most significant public health threats of our time^1, 2^. MDR Enterobacteriaceae are associated with significant morbidity and mortality in infected individuals, and an increasing number of reports describe extra-intestinal and invasive infections with these organisms in children^3-5^. Many studies show that beta-lactamase (*bla*) genes harbored by Enterobacteriaceae, e.g. extended-spectrum beta-lactamases (ESBLs or *bla*_ESBLs_) and carbapenemases (e.g., *bla*_KPCs_), are the major contributors to this growing problem^6, 7^. Strains carrying *bla* genes often harbor additional antibiotic resistance determinants residing on mobile genetic elements, e.g. plasmids and transposons, which are capable of rapid dissemination^1, 7^.

In the U.S. and worldwide, the CTX-M-type ESBL producing Enterobacteriaceae (CTX-M Ent) are the predominant ESBL-producing strains, with the clonal multi-locus sequence type (ST) 131 *E. coli* harboring CTX-M-15 (CTX-M-1 group) being the most commonly reported in both adult and pediatric studies^7, 8^. However, several other CTX-M types continue to circulate in Ent, and are equally concerning, highly resistant pathogens that can be acquired in the community^9^.

A notable example is the widespread CTX-M-9-group ESBL genes (e.g. *bla*_CTX-M-9_, *bla*_CTX-M-14_, *bla*_CTX-M-27_) which are the most common CTX-M genes circulating in Ent in some geographic regions^10^. A 2016 U.S. study of prevalence of CTX-M genes in Enterobacteriaceae found relative stability of *bla*_CTX-M-15_ in *E. coli*. Nevertheless, an increase in the prevalence of *bla*_CTX-M-14_ in *E. coli* occurred (25.6% and 20.4% in 2013 and 2014) compared to 2012 (16.0%)^11^. Despite this, much less is reported concerning the epidemiology of CTX-M-9-group infections. Unlike the clonal ST131 *E. coli* strains, CTX-M-9-group genes are associated with multiple strain types and horizontal gene transfer via mobile genetic elements likely plays an important role in their dissemination^12^. While studies of ESBL-Ent in U.S. children and internationally have reported infections due to CTX-M-9-group-producing Enterobacteriaceae (CTX-M-9-grp-Ent)^10, 13^, pediatric studies have not focused on the clinical and molecular epidemiology and impact of infections specifically due to CTX-M-9-grp-Ent.

In previous investigations, the genetic basis for beta-lactam resistance in Enterobacteriaceae isolates recovered from children cared for by multiple centers in the Chicago area was determined^13^. Subsequently, subgroups of children with infections due to similar resistance determinants, (e.g. plasmid-mediated fluoroquinolone, ESBL, and carbapenem resistance genes) were analyzed in order to determine genotypes, host factors, and exposures associated with specific MDR Enterobacteriaceae strains^13, 14^. A prior analysis of children with plasmid-mediated fluoroquinolone-resistant (PMFQR)-Ent infections that there are genetic and geospatial community links to MDR in our pediatric population was found ^15^. Here we report that CTX-M-9-grp-Ent infections in children are similarly linked to geographic location, and that acquisition of CTX-M-9-grp-Ent in children demonstrates environmental influences and originates in the community.

## METHODS

### Study Setting

Hospital A contains a 115-bed children’s hospital which includes pediatric and psychiatric wards, a mother-newborn infant unit, and cardiac, neonatal and pediatric intensive-care units (CICU, NICU and PICU), located within a tertiary care academic medical center. Hospital B is a free-standing children’s academic medical center comprised of 288 beds and provides quaternary care services, including bone marrow and organ transplantation. Hospital C is a 125-bed children’s hospital within an academic medical center, has newborn infant and general pediatrics wards, as well as a NICU and PICU. The participating centers are all located within metropolitan Chicago.

### Descriptive Study Design

#### Study Population

Patients included in this study were children aged 0 to 20.99 years who were identified because they possessed extended-spectrum cephalosporin (ceftriaxone, ceftazidime, cefotaxime) resistant (ESC-R) Enterobacteriaceae growing in clinical cultures and were suspected to harbor a *bla* gene conferring cephalosporin resistance by the clinical microbiology laboratories. Infections diagnosed between January 1, 2011 and December 31, 2016 were included in the analysis; only the first infection per patient was included. Study approval was obtained from the institutional review boards of the participating institutions and the need for informed consent was waived.

#### Antibiotic Susceptibility Testing in Enterobacteriaceae Isolates

The clinical microbiology laboratories of Hospitals A-C phenotypically analyzed ESC-R isolates by the Vitek 2 microbial identification system (*bioMérieux, Athens, GA)* or via the MicroScan WalkAway system (Beckman Coulter, Brea, CA). Following Clinical and Laboratory Standards Institute (CLSI) guidelines, one or more of the following antimicrobials: aztreonam, ceftriaxone, ceftazidime, cefotaxime or cefpodoxime were used to screen for ESBL production^16^. Techniques used to confirm ESBL production included automated instruments or disk diffusion assays (BBL; Becton, Dickinson and Company, Sparks, MD) to measure cefotaxime and ceftazidime susceptibility in the presence and absence of clavulanic acid. An increase in measure of a disk zone diameter of > 5 mm or a 4-fold reduction in the minimum inhibitory concentration (MIC) of cefotaxime and ceftazidime in the presence of clavulanic acid served as confirmation of the ESBL phenotype^16^.

### Molecular Analysis

#### Beta-Lactam Resistance Determinants

Genomic DNA from Enterobacteriaceae isolates was purified from isolates confirmed with an ESBL phenotype (DNeasy blood and tissue kit, Qiagen, Inc. Valencia, CA). DNA Microarray (Check-MDR CT101 and CT103XL; Check-Points, Wageningen, Netherlands) and polymerase chain reaction (PCR) was performed to assess and confirm the presence of *bla* genes in isolates as previously described and according to manufacturer protocol^13, 17^. Isolates found to be positive solely for beta-lactamase genes belonging to CTX-M-9-grp were included in the analysis.

#### Nomenclature and Characterization

A well-established multiplex PCR-based method was used to assign *E. coli* to one of four phylogenetic groups (A, B1, B2 and D)^18^. Multi-locus sequence typing (MLST) [Pasteur website (http://www.pasteur.fr/recherche/genopole/PF8/mlst/)] and DNA sequencing identified sequence types and PCR distinguished *bla*_CTX-M-9-grp_ alleles in the ESBL-producing strains of *E. coli* and *Klebsiella* species as previously described^13, 19, 20^. Plasmids were typed based on incompatibility groups corresponding to the nomenclature assigned by Carattoli *et al*.^21^

#### Analytic Study Design

A retrospective case–control study design was used to assess factors associated with infection due to isolates in which we had detected a *bla*_CTX-M-9-grp_ gene. Children serving as control subjects were identified using hospital electronic laboratory records (ELRs) and were matched 3:1 to the cases by age range, hospital, and specimen source, a design previously described^22, 23^. Only children diagnosed with clinical infections due to bacteria susceptible to extended-spectrum cephalosporins (ESC-S) were included, as determined by study investigator case review and/or using standard criteria defined by the CDC National Healthcare Safety Network^24^.

#### Covariates

We analyzed several variables as potential factors associated with CTX-M-9-grp-Ent infection based on known associations for ESBL-producing Enterobacteriaceae acquisition in adults and children including (1) demographics (age, gender, race/ethnicity); (2) comorbid conditions (as defined by ICD-9/ICD-10 codes); (3) recent inpatient and outpatient healthcare exposures, including hospitalization and/or procedures in the previous 30 days; (4) antibiotic exposures in the 40 days prior to culture^22^; (5) presence, number, and type of invasive medical devices; and (6) patient residence in the Chicago area as assessed by dividing the metropolitan area into 7 regions using zip code level data, which included Chicago proper and its suburban areas (i.e. North side and North Suburbs, South side and South Suburbs, etc.). An eighth region was included for patients residing in other states or other parts of Illinois.

#### Statistical Analysis

We examined case and control groups for differences using parametric or non-parametric tests as appropriate for categorical and continuous variables; a p-value of ≤0.05 was considered statistically significant unless otherwise specified. Bivariate analysis was performed and variables with p<0.1 were included in multivariable analysis. To assess the multivariable relationship between the covariates and the groups, stepwise multiple logistic regression was employed. The simplest model was included in the final multivariable logistic regression with significant covariates (p<0.05) from the stepwise selection process, with CTX-M-9-grp-Ent infection as the outcome variable. The simplest model was chosen based on a relatively small sample size and the effect of variables in the model. All analyses were performed in SAS 9.4 (SAS Institute, Cary, NC, USA).

## RESULTS

Of 356 pediatric Enterobacteriaceae isolates found to harbor 350 *bla* genes between 2011 - 2016, 44 (12.4%) were positive by DNA microarray and PCR amplification solely for *bla*_CTX-M-9-grp_ genes. Most (91%) CTX-M-9-grp-Ent infections, and a significantly lesser fraction (58%) of ESC-S infections, were *E. coli*, p=0.003 (Table 1). Urine was the most common source of CTX-M-9-grp-Ent (70%); differences in the source of infection between groups were not observed. The antimicrobial susceptibility testing results of CTX-M-9-grp-Ent are summarized in Table 2. The highest retained susceptibility was to amikacin, carbapenems and piperacillin-tazobactam. Multi-drug resistance (resistance to ≤3 antibiotic classes) was found in 77% of case isolates.

| Characteristics of CTX M-9-grp extended spectrum beta-lactamase (ESBL) producing Enterbacteriaceae and ESC<sup>a</sup> sensitive Enterobacteriaceae (ESC-S) |  |  |
| --- | --- | --- |
|  | CTX M-9-grp ESBL<br>(n=44) | ESC-S (n=135) |
| **Species n (%)** |  |  |
| *Klebsiella pneumoniae* | 3 (7) | 24 (18) |
| *Escherichia coli* | 40 (91) | 79 (58) |
| *Enterobacter cloacae* | 1 (2) | 22 (16) |
| *Other Enterobacteriaceae* | 0 (0) | 10 (7) |
| **Source n (%)** |  |  |
| Urine | 31 (70) | 99 (73) |
| Respiratory | 7 (16) | 20 (15) |
| Blood | 4 (9) | 8 (6) |
| Other<sup>b</sup> | 2 (5) | 8 (6) |

<sup>a</sup> ESC, Extended Spectrum Cephalosporin
<sup>b</sup>Other includes wound, abscess, abdominal and peritoneal sources

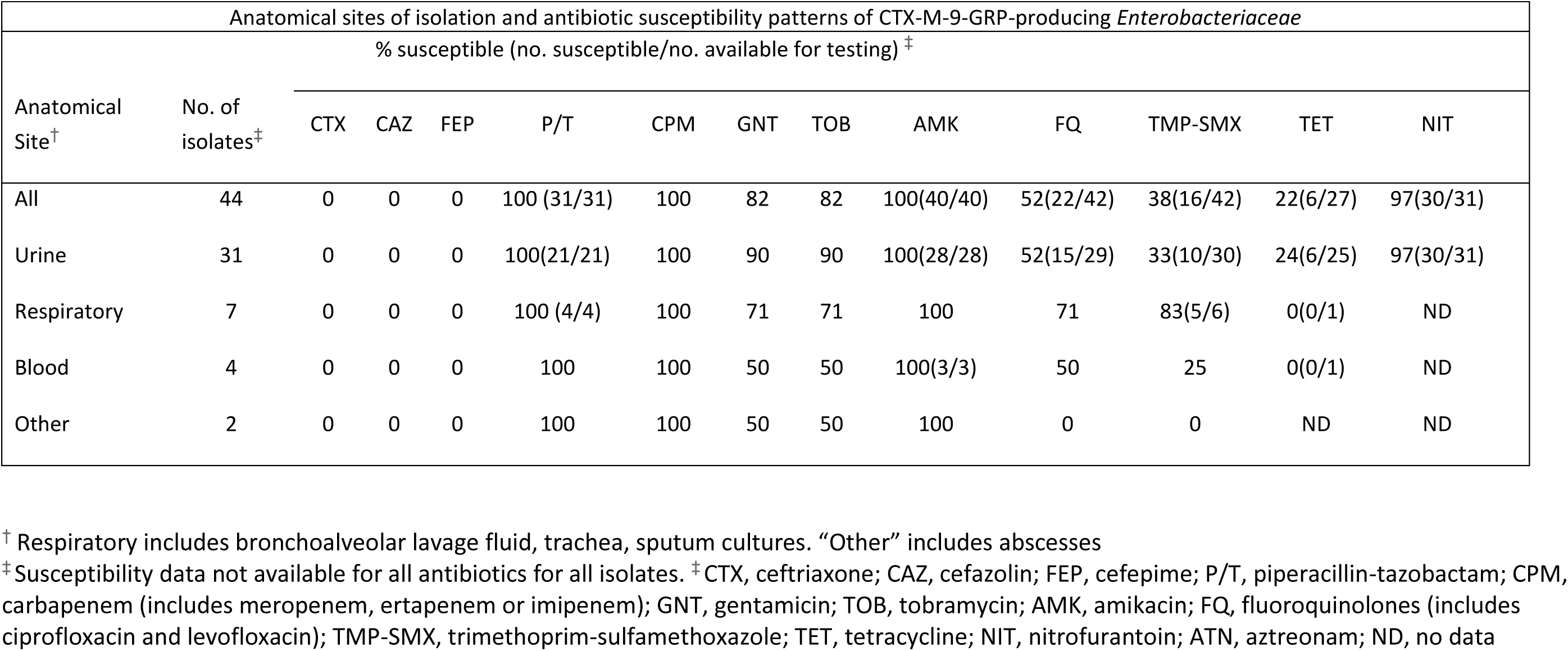

### Molecular analysis

Of *E. coli*, 60% belonged to phylogenetic group D; the rest belonged to B2. Both are virulent phylogroups associated with extra-intestinal pathogenic *E. coli* infections. Of CTX-M-9-group, the most common subtypes were CTX-M-14 (61%) and CTX-M-27 (34%). MLST revealed diverse strain types. Of the 10 ST types associated with 40 CTX-M-9 *E. coli*, 25% were ST8; ST43/ST131 and ST506 were the next most common (20% and 15%, respectively). Two novel *E. coli* strain types were also discovered to carry *bla*_CTX-M-9-grp_ (data not shown).

### Analysis of Factors Associated with CTX-M-9 Ent Infection

The 44 cases of CTX-M-9-grp-Ent infection were matched to 135 ESC-S controls. On bivariate analysis (Table 3) factors associated with higher likelihood of CTX-M-9-grp-Ent infection compared to ESC-S controls included: infection with *E. coli*, diagnosis in outpatient clinic, absence of recent health care, residence in the South or Southwest regions (comprising south and southwest Chicago and associated suburbs), and race “other” (non-white, non-black, non-Hispanic). Children with CTX-M-9-grp-Ent infections were less likely than controls to have infection with *Enterobacter* sp., diagnosis in the inpatient, non-ICU setting (i.e. general pediatric wards), residence in Northwest region (comprised of Northwestern Chicago and the Northwestern suburbs), presence of a central venous line, respiratory comorbidities, and recent outpatient health care (including outpatient procedures).

**TABLE 3.** BIVARIATE ANALYSIS OF DEMOGRAPHICS AND FACTORS ASSOCIATED WITH CTX-M-9-GRP ENTEROBACTERIACEAE INFECTION

| Characteristic<sup>a</sup> | CTX-M-9-GRP Ent<br>Infection | ESC-S Ent Infection<sup>b</sup> | p value |
| --- | --- | --- | --- |
| Patient | n=44 | n=135 |  |
| Age | 4.14 (0-20.8) | 4.52 (0-20.9) | 0.083 |
| Female | 29 (34.1) | 93 (31.1) | 0.713 |
| Race, White | 15 (34.1) | 40 (29.6) | 0.58 |
| Race, Black | 3 (6.8) | 29 (21.5) | 0.04 |
| Race, Hispanic | 15 (34.1) | 59 (43.7) | 0.26 |
| Race, Non-White,<br>Non-Black, Non-Hispanic | 11 (25) | 7 (5.2) | 0.0001 |
| Location at Diagnosis |  |  | <0.0001 |
| Inpatient, non ICU | 7 (15.9) | 43 (31.9) | 0.04 |
| Outpatient Clinic | 16 (36.4) | 10 (7.4) | <0.001 |
| Emergency Room | 9 (20.5) | 24 (17.8) | 0.69 |
| Pediatric ICU | 9 (20.5) | 40 (29.6) | 0.24 |
| Neonatal ICU | 3 (6.8) | 18 (13.3) | 0.11 |
| Recent CTX<sup>e</sup> | 9 (15.9) | 33 (24.4) | 0.24 |
| Recent FQ<sup>f</sup> | 3 (6.8) | 4 (3.0) | 0.37 |
| Recent TMP/SMX<sup>g</sup> | 5 (11.4) | 18 (13.5) | 0.71 |
| Region of Residence<sup>c</sup> |  |  | 0.009 |
| Recent Health Care |  |  | 0.008 |
| Inpatient Care | 11 (25) | 34 (25.2) | 0.98 |
| Outpatient Care<sup>d</sup> | 10 (22.7) | 62 (45.9) | 0.006 |
| No Recent Care | 23 (52.3) | 39 (28.9) | 0.005 |
| Foreign body Present | 19 (43.2) | 77 (57.0) | 0.11 |
| Multiple Foreign<br>Bodies Present | 13 (29.6) | 50 (37.0) | 0.37 |
| Central venous line | 8 (18.2) | 45 (33.3) | 0.025 |
| Comorbidities |  |  |  |
| Cardiovascular | 7 (15.9) | 29 (21.5) | 0.42 |
| Endocrine | 2 (4.6) | 9 (6.8) | 0.73 |
| Gastrointestinal | 12 (27.3) | 33 (24.4) | 0.71 |
| Genitourinary | 14 (26.4) | 37 (28.2) | 0.80 |
| Hematology | 6 (13.6) | 22 (16.3) | 0.67 |
| Neurology | 14 (31.8) | 48 (35.6) | 0.65 |
| Respiratory | 8 (18.2) | 47 (34.8) | 0.04 |
| Renal | 13 (29.6) | 42 ( 31.1) | 0.85 |
| Multiple<br>Comorbidities | 18 (40.9) | 63 (46.7) | 0.51 |

<sup>a</sup> Values represent n (%) unless otherwise indicated.
<sup>b</sup> ESC-S Ent Infection were children with infections due to Enterobacteriaceae sensitive to extended spectrum cephalosporin antibiotics.
<sup>c</sup> Region of residence includes city region and neighboring suburbs
<sup>d</sup> Outpatient care includes care outside of routine child care visits and outpatient procedures.
<sup>e</sup> Abbreviation CTX includes extended-spectrum cephalosporins (ceftriaxone, ceftazidime, cefotaxime, cefepime)
<sup>f</sup> FQ, Fluoroquinolones (ciprofloxacin and levofloxacin)
<sup>g</sup> TMP-SMX, Trimethoprim-Sulfamethoxazole.

During model building stages, we did not find evidence of significant effect modification or significant confounding; therefore, additional covariates were not added back to the final model after the stepwise selection process was completed, and the simplest model with significant covariates was included in the final regression model.

In the multivariable regression analysis (Table 4), factors associated with CTX-M-9-grp-Ent infection included having infection due to *E. coli* (OR 7.0; 95% CI 2.2, 22.2; p<0.001); being of a race or ethnicity “other” (OR 6.5; 95% CI 1.7, 24.3; p=0.002); and being diagnosed in the outpatient clinic setting (OR 4.5; 95% CI 1.7, 11.9; p=0.003).

**TABLE 4.** MULTIVARIABLE ANALYSIS OF FACTORS ASSOCIATED WITH CTX-M-9-GRP ENTEROBACTERIACEAE INFECTIONS IN CHILDREN

| Associated Factor with CTX-M-9-grp Ent infection<sup>a</sup> | OR | 95% CI | p value |
| --- | --- | --- | --- |
| Infection with *E. coli* | 7.0 | 2.2, 22.2 | 0.001 |
| Home Residence in South Region (South Chicago and South Suburbs) | 6.7 | 1.2, 37.7 | 0.02 |
| Race not white, black or Hispanic | 6.5 | 1.70, 24.3 | 0.002 |
| Outpatient Clinic Location at time of infection diagnosis | 4.5 | 1.69, 11.9 | 0.003 |
| Home Residence in Northwest Region (NW Chicago and NW Suburbs) | 0.19 | 0.04, 0.84 | 0.02 |

<sup>a</sup>Ent, Enterobacteriaceae
<sup>b</sup>Reference Region, West Region.

Remarkably, among children with Enterobacteriaceae infections, those residing in the South region of Chicago demonstrated more than six times the odds of having a CTX-M-9-grp-Ent infection compared to those living in the reference West Chicago region (OR 6.7; 95% CI 1.2, 37.7; p=0.02) after controlling for race, infecting organism, and healthcare setting. In contrast, for children who resided in the Northwest region, there was an 81% decrease in the odds of CTX-M-9-grp-Ent infection compared to those residing in the reference West Chicago region (OR 0.19; 95% CI 0.04, 0.84; p=0.02). Residence in the Southwest region (the neighboring region to the South region) was also significant for increased likelihood of CTX-M-9-grp infection on bivariate analysis (OR 2.8; 95% CI 1.02, 7.6; p=0.04).

## DISCUSSION

Our analysis is the first multi-centered study to assess the clinical and molecular epidemiology of CTX-M-9 type ESBL-producing Enterobacteriaceae infections in children located in an urban setting. We found that *E. coli* was the most common infecting organism associated with *bla*_CTX-M-9-grp_. The CTX-M-9 group are thought to be the second most commonly community-acquired ESBL genes in Enterobacteriaceae worldwide, and, similar to the high-risk clonal ST131 CTX-M-15 producing *E. coli*, are reported in people without significant healthcare exposure and are associated with multiple antibiotic gene cassettes leading to MDR^3, 9^. However, unlike the association of CTX-M-1 group with ST131, we found that *bla*_CTX-M-9-grp_ genes are carried by multiple strain types, including the identification of novel STs associated with pediatric infection.

Because of multiple associated strain types carrying *bla*_CTX-M-9-grp_ genes, we hypothesized that there is significant horizontal gene transfer occurring between genera. Plasmid replicon typing of isolates also supports this hypothesis, as multiple plasmids of different types were found present in isolates from the “high-risk” South region, with Inc F plasmids (FII, FIA, FIB) being the most common (data not shown). This observation is concerning as children once colonized with MDR Enterobacteriaceae can remain colonized for months to years, and young children (median age 4.1 years) were carrying organisms with multiple plasmids, capable of rapid dissemination and spread^13, 25^. Additionally, we found novel ST types carrying *bla*_CTX-M-9-grp_ genes. This portends a troublesome epidemiology suggesting the acquisition of these genes may be occurring in commensal or previously non-pathogenic strains of Enterobacteriaceae.

In our study, most (77%) CTX-M-9-grp-Ent isolates were phenotypically MDR. While the highest retained antibiotic susceptibility included piperacillin-tazobactam, clinicians should be cautioned in its use for serious, invasive infections (e.g. bacteremia). Recent randomized clinical trials have demonstrated inferiority of piperacillin-tazobactam to carbapenem therapy when used to treat ESC-R Ent bloodstream infections (including ESBL-producers) and describe worse patient outcomes associated with piperacillin-tazobactam use as definitive therapy^26^. Of additional concern, susceptibility to oral agents was significantly reduced leaving few oral options for step-down therapy for children (Table 2). The sole oral exception was nitrofurantoin (97% sensitivity) which can be used to treat uncomplicated lower urinary tract infection but should not be used for complicated, invasive infections or acute pyelonephritis, due to the inability of the drug to attain therapeutic levels in the renal parenchyma and bloodstream^27^.

The residential differences for children infected with CTX-M-9-grp-Ent compared to children infected with antibiotic sensitive strains was striking. We found a substantial increase in odds of infection with the resistant strains in south Chicago region, and a significant decrease in odds of infection in the northwest Chicago region. Our study included 3 major pediatric centers, none of which are located in the “high-risk” South region, yet all three centers diagnosed and treated patients with CTX-M-9-grp-Ent infections from these high-risk regions. The reservoirs associated with acquisition of CTX-M-9-grp-Ent in these regions are currently undefined.

Of additional concern, in a separate study of plasmid-mediated fluoroquinolone resistant (PMFQR) Ent infections in children between 2011-2014, we found a substantial increase in odds of PMFQR Ent infection in the Southwest region, while in the Downtown region (comprising the downtown Chicago area, near North side, Chicago loop, and North Chicago) there was a significant decrease in the likelihood of PMFQR Ent infection^15^. These PMFQR Ent “high-risk” and “low-risk” regions neighbor the “high-risk” South region and “low-risk” Northwest region in the current study. The Southwest region was found to be associated with an increase in the odds of CTX-M-9-grp-Ent infection on bivariate analysis in this study, though this did not reach statistical significance on multivariable analysis once controlling for other variables.

Furthermore, we found that children with CTX-M-9-grp-Ent infection were significantly more likely to present in the outpatient clinic setting compared to those with antibiotic susceptible Ent infections. This is surprising and striking as most infections caused by MDROs in the U.S. are linked to healthcare settings^28^. While the majority of infections were associated with virulent extra-intestinal-pathogenic *E. coli* phylogenetic groups (B and D), these data suggest that CTX-M-9-grp-Ent infections in children are being acquired in the community and may be less severe at presentation as children presented more often in outpatient clinics. This is in stark contrast to the epidemiology of MDR Enterobacteriaceae infections reported in adults in Chicago, where acquisition has been strongly associated with residence in long-term care facilities and the interfacility transfer of patients^29^.

The higher likelihood of CTX-M-9-grp-Ent infection in those of non-white, non-black, and non-Hispanic race was an additional association discovered on multivariable analysis. This risk was statistically independent of residence, and only one of the children located in the “high risk” south region was of race “other”, supporting the independence of the two risk factors for infection. We were unable to gather further data on “other” race or ethnicity due to the retrospective nature of the study, and we did not have travel data for the majority of children. It is well described that travel to certain countries can be associated with high rates of ESBL Ent acquisition, particularly in South and Southeast Asia^30^. It is also well described that there is an increased risk of colonization when residing in a household with a family member with history of ESBL Ent or who acquired such a strain during travel to an endemic region^31^.

Environmental influences originating in the community could include higher risks in certain populations due to specific exposures to foods, animals, water sources, fertilizer, soil, and vegetation^32^. We did not have data on companion pets for the majority of children and were unable to examine this risk factor; though pets may play a significant role in human acquisition of antibiotic resistant bacteria^33^. Lastly, we did not find overall differences in a general comparison of socioeconomic status of the “high risk” south region and neighboring regions such as the southwest and west regions using regional zip codes and Illinois census data.

We recognize the limitations of our study. This was a retrospective study of children with Enterobacteriaceae infections in a single metropolitan area which may affect generalizability to other regions. While selection bias may result from small sample sizes, the pooling of multicentered data from institutions of differing types in the 3rd largest U.S. metropolitan area and serving diverse populations potentially lessens this bias. Moreover, our sample size is consistent with the overall low prevalence of these MDR Enterobacteriaceae children in the majority of the U.S. (1-3%) including Chicago and the Midwest region^34, 35^; however, it should be noted that national trends indicate an increase in the incidence and prevalence of pediatric MDR Enterobacteriaceae over the last decade, an emerging threat that needs further assessment^5, 36^.

In conclusion, we describe, for the first time, community origins of CTX-M-9-grp Enterobacteriaceae in children, and the impact of residence on infection in children located in the same geographic area. The reservoirs associated with CTX-M-9-grp-Ent infections remain undefined. Future studies must focus on environmental influences associated with regional acquisition. “Silent dissemination” of community-acquired MDR Enterobacteriaceae is likely occurring in children outside of healthcare settings. Dedicated programs on a local, national and global scale must focus on children and halting of spread of these dangerous pathogens.

## Acknowledgements

We gratefully acknowledge the contribution of the late Dr. Paul Schreckenberger to this work. We thank the microbiology laboratories of the participating institutions for providing isolates for this study. We thank Laura Rojas-Coy for contributions in the CTX-M primer design for PCR amplification. We thank Kendrick Reme and Lynika Strozier of the Logan Laboratory and Pamela Hagen, Jane Stevens, Joyce Houlihan, Kathleen McKinley, Violeta Rekasiu, Cindy Bethel, and Donna Carter of participating institutions for collection, shipping, and cultivation of organisms. We thank the team of curators of the Institut Pasteur MLST and whole-genome MLST databases for curating the data and making them publicly available at http://bigsdb.web.pasteur.fr/. We report no conflicts of interest relevant to this study. The content is solely the responsibility of the authors and does not necessarily represent the official views of the National Institutes of Health or the Department of Veterans Affairs.

## References

1. Logan LK, Weinstein RA. The Epidemiology of Carbapenem-Resistant Enterobacteriaceae: The Impact and Evolution of a Global Menace. J Infect Dis. 2017;215:S28–S36.

2. Medernach RL, Logan LK. The Growing Threat of Antibiotic Resistance in Children. Infect Dis Clin North Am. 2018;32:1–17.

3. Lukac PJ, Bonomo RA, Logan LK. Extended-spectrum beta-lactamase-producing Enterobacteriaceae in children: old foe, emerging threat. Clinical Infectious Diseases. 2015;60:1389–1397.

4. Logan LK, Renschler JP, Gandra S, et al. Carbapenem-Resistant Enterobacteriaceae in Children, United States, 1999–2012. Emerging Infectious Diseases. 2015;21:2014–2021.

5. Logan LK, Braykov NP, Weinstein RA, Laxminarayan R, CDC Epicenters Prevention Program. Extended-Spectrum beta-Lactamase-Producing and Third-Generation Cephalosporin-Resistant Enterobacteriaceae in Children: Trends in the United States, 1999–2011. Journal of the Pediatric Infectious Diseases Societ. 2014;3:320–328.

6. Munoz-Price LS, Poirel L, Bonomo RA, et al. Clinical epidemiology of the global expansion of Klebsiella pneumoniae carbapenemases. The Lancet infectious diseases. 2013;13:785–796.

7. Price LB, Johnson JR, Aziz M, et al. The epidemic of extended-spectrum-beta-lactamase-producing Escherichia coli ST131 is driven by a single highly pathogenic subclone, H30-Rx. MBio. 2013;4:e00377–13.

8. Miles-Jay A, Weissman SJ, Adler AL, et al. Epidemiology and Antimicrobial Resistance Characteristics of the Sequence Type 131-H30 Subclone Among Extraintestinal Escherichia coli Collected From US Children. Clinical Infectious Diseases. 2017;66:411–419.

9. Cantón R, González-Alba JM, Galán JC. CTX-M enzymes: origin and diffusion. Frontiers in microbiology. 2012;3:110.

10. Merida-Vieyra J, De Colsa A, Castañeda YC, Barbosa PA, Andrade AA. First Report of Group CTX-M-9 Extended Spectrum Beta-Lactamases in Escherichia coli Isolates from Pediatric Patients in Mexico. Plos one. 2016;11:e0168608.

11. Castanheira M, Mendes RE, Jones RN, Sader HS. Changes in the Frequencies of beta-Lactamase Genes among Enterobacteriaceae Isolates in U.S. Hospitals, 2012 to 2014: Activity of Ceftazidime-Avibactam Tested against beta-Lactamase-Producing Isolates. Antimicrob Agents Chemother. 2016;60:4770–4777.

12. Oteo J, Diestra K, Juan C, et al. Extended-spectrum β-lactamase-producing Escherichia coli in Spain belong to a large variety of multilocus sequence typing types, including ST10 complex/A, ST23 complex/A and ST131/B2. Int J Antimicrob Agents. 2009;34:173–176.

13. Logan LK, Hujer AM, Marshall SH, et al. Analysis of beta-Lactamase Resistance Determinants in Enterobacteriaceae from Chicago Children: a Multicenter Survey. Antimicrobial Agents & Chemotherapy. 2016;60:3462–3469.

14. Logan LK, Nguyen DC, Scaggs-Huang FA, et al. A multi-centered case-case-control study of factors associated with Klebsiella pneumoniae carbapenemase (KPC)-producing Enterobacteriaceae (KPC-CRE) infections in children and young adults. Pediatric Infectious Disease Journal. 2019;38.

15. Logan LK, Medernach RL, Rispens JR, et al. Community origins and regional differences in plasmid-mediated fluoroquinolone resistant Enterobacteriaceae infections in children. bioRxiv. 2018:301457.

16. Clinical and Laboratory Standards Institute. Performance Standards for Antimicrobial Susceptibility Testing: Twentieth Informational Supplement (June 2010 Update). 2010:December 29, 2011.

17. Powell EA, Haslam D, Mortensen JE. Performance of the check-points check-MDR CT103XL assay utilizing the CDC/FDA antimicrobial resistance isolate bank. Diagnostic Microbiology & Infectious Disease. 2017;88:219–221.

18. Bingen-Bidois M, Clermont O, Bonacorsi S, et al. Phylogenetic analysis and prevalence of urosepsis strains of Escherichia coli bearing pathogenicity island-like domains. Infect Immun. 2002;70:3216–3226.

19. Diancourt L, Passet V, Verhoef J, Grimont PA, Brisse S. Multilocus sequence typing of Klebsiella pneumoniae nosocomial isolates. J Clin Microbiol. 2005;43:4178–4182.

20. Jaureguy F, Landraud L, Passet V, et al. Phylogenetic and genomic diversity of human bacteremic Escherichia coli strains. BMC Genomics. 2008;9:560-2164-9-560.

21. Carattoli A, Bertini A, Villa L, Falbo V, Hopkins KL, Threlfall EJ. Identification of plasmids by PCR-based replicon typing. J Microbiol Methods. 2005;63:219–228.

22. Logan LK, Meltzer LA, McAuley JB, et al. Extended-Spectrum beta-Lactamase-Producing Enterobacteriaceae Infections in Children: A Two-Center Case-Case-Control Study of Risk Factors and Outcomes in Chicago, Illinois. Journal of the Pediatric Infectious Diseases Societ. 2014;3:312–319.

23. Medernach RL, Rispens JR, Marshall SH, et al. Resistance Mechanisms and Factors associated with Plasmid-Mediated Fluoroquinolone Resistant (PMFQR) Enterobacteriaceae (Ent) Infections in Children. ASM Microbe. 2017.

24. Centers for Disease Control and Prevention (CDC). The National Healthcare Safety Network (NHSN) Patient Safety Component Manual. 2017.

25. Zerr DM, Qin X, Oron AP, et al. Pediatric infection and intestinal carriage due to extended-spectrum-cephalosporin-resistant Enterobacteriaceae. Antimicrob Agents Chemother. 2014;58:3997–4004.

26. Harris PN, Peleg AY, Iredell J, et al. Meropenem versus piperacillin-tazobactam for definitive treatment of bloodstream infections due to ceftriaxone non-susceptible Escherichia coli and Klebsiella spp (the MERINO trial): study protocol for a randomised controlled trial. Trials. 2015;16:24.

27. Beetz R, Westenfelder M. Antimicrobial therapy of urinary tract infections in children. Int J Antimicrob Agents. 2011;38:42–50.

28. Sievert DM, Ricks P, Edwards JR, et al. Antimicrobial-resistant pathogens associated with healthcare-associated infections summary of data reported to the National Healthcare Safety

29. Network at the Centers for Disease Control and Prevention, 2009–2010. Infection Control & Hospital Epidemiology. 2013;34:1–14.

30. Snitkin ES, Won S, Pirani A, et al. Integrated genomic and interfacility patient-transfer data reveal the transmission pathways of multidrug-resistant Klebsiella pneumoniae in a regional outbreak. Sci Transl Med. 2017;9: 10.1126/scitranslmed.aan0093.

31. Kuenzli E, Jaeger VK, Frei R, et al. High colonization rates of extended-spectrum ß-lactamase (ESBL)-producing Escherichia coli in Swiss Travellers to South Asia–a prospective observational multicentre cohort study looking at epidemiology, microbiology and risk factors. BMC infectious diseases. 2014;14:528.

32. Haverkate MR, Platteel TN, Fluit A, et al. Quantifying within-household transmission of extended-spectrum β-lactamase-producing bacteria. Clinical Microbiology and Infection. 2017;23:46. e1-46. e7.

33. Silbergeld EK, Graham J, Price LB. Industrial food animal production, antimicrobial resistance, and human health. Annu Rev Public Health. 2008;29:151–169.

34. Grasselli E, François P, Gutacker M, et al. Evidence of horizontal gene transfer between human and animal commensal Escherichia coli strains identified by microarray. FEMS Immunology & Medical Microbiology. 2008;53:351–358.

35. Logan LK, Meltzer LA, McAuley JB, et al. Extended-Spectrum β-Lactamase–Producing Enterobacteriaceae Infections in Children: A Two-Center Case–Case–Control Study of Risk Factors and Outcomes in Chicago, Illinois. Journal of the Pediatric Infectious Diseases Society. 2014;3:312–319.

36. Zerr DM, Weissman SJ, Zhou C, et al. The Molecular and Clinical Epidemiology of Extended-Spectrum Cephalosporin–and Carbapenem-Resistant Enterobacteriaceae at 4 US Pediatric Hospitals. Journal of the Pediatric Infectious Diseases Society. 2017:piw076.

37. Meropol SB, Haupt AA, Debanne SM. Incidence and outcomes of infections caused by multidrug-resistant enterobacteriaceae in children, 2007–2015. Journal of the Pediatric Infectious Diseases Society. 2017;7:36–45.

